# Hydrological dynamics drive the transition of antibiotic resistance genes between particle-attached and free-living lifestyles in a deep freshwater reservoir

**DOI:** 10.1101/2022.06.11.495731

**Authors:** Bob Adyari, Liyuan Hou, Lanping Zhang, Nengwang Chen, Feng Ju, Longji Zhu, Chang-Ping Yu, Anyi Hu

## Abstract

Despite the growing awareness of antibiotic resistance genes (ARGs) spreading in the environment, there is a knowledge gap on the fate and transport of ARGs in particle-attached (PA) and free-living (FL) lifestyles in deep freshwater ecosystems experiencing seasonal hydrological changes. Here, we examined the ARG profiles using high-throughput quantitative PCR in PA and FL lifestyles at four seasons representing two hydrological seasons (i.e., vertical mixing and thermal stratification) in the Shuikou Reservoir (SR), Southern China. The results indicated that seasonal hydrological dynamics were critical for influencing ARGs in PA and FL fractions, and the transition of ARGs between the two lifestyles. Although both PA and FL ARG profiles were likely to be shaped by horizontal gene transfer, PA and FL ARGs had different responses to the changes in physico-chemicals (e.g., nutrients and dissolved oxygen) caused by seasonal hydrological dynamics. The particle-associated niche (PAN) index revealed that there were 94 non-conservative ARGs (i.e., no preferences for PA and FL), 23 conservative ARGs that preferred PA lifestyle, and 16 conservative ARGs for FL lifestyle. A sharp decline in the number of conservative ARGs in stratified seasons suggests a hydrological dynamics-dependent transition of ARGs between two lifestyles. Remarkably, the conservative ARGs (in PA or FL lifestyle) were more closely related to bacterial OTUs in their preferred lifestyle compared to their counterpart lifestyle, suggesting a lifestyle-dependent ARG enrichment. Altogether, these findings enhance our understanding of the role of seasonal hydrological changes in the dissemination of ARGs in different size fractions in deep aquatic ecosystems.

## Introduction

Prevalent antibiotic use has enriched the antibiotic resistance bacteria (ARB) and enhanced the spread of antibiotic resistance genes (ARGs) in human/animal guts ^1,2^. The discharge of municipal and livestock wastewater, together with agricultural run-off can lead to an increase in ARG pollution levels in receiving water bodies, including rivers ^3^ and reservoirs ^4^ through horizontal gene transfer (HGT) and vertical gene transfer (VGT). The HGT is mainly mediated by mobile genetic elements (MGEs) and has been reported as the main mechanism for the spread of ARGs in aquatic ecosystems ^3–5^. However, some studies found that VGT could be the essential mechanism shaping the ARG profiles of microbial communities under certain scenarios ^6,7^. The enhanced ARG levels in the environment may pose potential harm to the health of aquatic ecosystems and cause an increased risk of disease transmission ^8,9^. Since freshwater reservoirs are key drinking water resources, a comprehensive understanding of ARG profiles and mechanisms associated with their dissemination in the reservoir is of great importance for making suitable management policies to control the ARGs and reduce the health risk.

Recent studies reported the occurrence of ARGs in freshwater reservoirs ^4,10^. One study found that ARGs had seasonal variations in reservoir ^4^, where abiotic (e.g., antibiotics and nutrients) and biotic (e.g., MGEs and cyanobacterial blooms) factors played essential roles in influencing the abundance and composition of ARGs ^10^. However, there is a knowledge gap regarding the seasonal-vertical variations of ARGs, their dissemination mechanisms, and the influencing factors in deep reservoirs experiencing vertical mixing and thermal stratification. Unlike their shallow counterpart, deep reservoirs usually undergo a complete vertical mixing from winter to spring and a thermal stratification from summer to fall ^11^. Due to the lack of vertical mixing of water columns in stratified seasons (e.g., summer and fall), temperature and dissolved oxygen (DO) significantly decrease with depth ^12^. This phenomenon can subsequently result in dramatic changes in the composition and function of microbial communities between upper and deeper layers ^13,14^. Therefore, it would be expected that ARG profiles may have similar depth-related variations, especially during the stratified seasons.

Planktonic microorganisms in aquatic environments can be empirically classified into particle-attached (PA, > 3 µm) and free-living (FL, between 3 µm and 0.22 µm) fractions based on their lifestyle preference ^15,16^. The suspended particles have been considered the hotspot of microbial diversity, biomass, and activity due to the high adsorption of organic and inorganic nutrients on the particle surface ^15,17^. Numerous studies have demonstrated that PA and FL fractions tended to have distinct microbial community compositions in diverse aquatic ecosystems ^16,18,19^. This is due to that most microbial phyla/classes have different lifestyle preferences (i.e., PA or FL lifestyle) ^16^. Previous studies have shown that PA and FL bacterial communities exhibit different responses to various environmental parameters ^14,20^. Likely, Guo et al. (2018) have found that ARGs in PA and FL lifestyles showed different responses (e.g., abundances, diversities, and compositions) to the cyanobacterial blooming event in a subtropical reservoir ^10^. These findings suggest that different abiotic and biotic factors may be responsible for regulating the dynamics of ARGs in these two lifestyles. Moreover, a few reports suggested that some microbial taxa can change their lifestyle preference (i.e., from PA to FL lifestyle or vice versa) at temporal-vertical scales to adapt to changes in environmental conditions ^18,19,21^. For instance, Mestre et al. (2017) found that *Rhodobacterales* (Alphaproteobacteria), SAR11 (Alphaproteobacteria), and Sh765B (Deltaproteobacteria) showed non-conservative preferences along the water column in the northwestern Mediterranean Sea. Another study revealed that SAR11 could change its lifestyle from FL to PA during the cold season ^22^. Despite this, little information is available regarding the lifestyle preference of ARGs for PA and FL fractions in aquatic ecosystems. More questions need to be answered: Does the majority of ARGs prefer a non-conservative lifestyle or a conservative lifestyle? Moreover, what are the environmental factors and the associated dissemination mechanisms for controlling this lifestyle pattern?

In this study, we used high-throughput quantitative PCR (HT-qPCR) to comprehensively profile the ARGs in a deep reservoir (Shuikou Reservoir, SR), Southern China. A total of 285 ARGs and 10 MGEs which cover the most common types and mechanisms of resistance were examined in both FL and PA fractions along a water depth gradient during four seasons. We hypothesized that: i) the abiotic factors (e.g., temperature, DO, etc.) related to changes in seasonal-hydrological conditions (i.e., vertical mixing and thermal stratification) would influence the ARG profiles in PA and FL fractions as well as their transition between PA and FL fractions; ii) MGEs may not only play an important role in the spread of ARGs in PA and FL fractions, but also in the transition between PA and FL fraction. PA rather than FL fraction may be the main HGT hotspot; and iii) PA and FL fractions may have distinct co-occurrence network patterns consisting of non-conservative and conservative ARGs. Particularly, more associations between ARGs and bacterial taxa (as potential ARG hosts) with the same lifestyle preference (i.e., FL or PA lifestyle) are highly expected. This study will promote our understanding of the role of seasonal mixing and thermal stratification in the spread of ARGs in PA and FL fractions in deep aquatic ecosystems.

## Materials and methods

### 2.1 Study area, sample collection, and processing

Min River is the largest river in Fujian Province, China, with a mainstream length of 562 km. It is the main water source of economic and cultural activities (e.g., agricultural, residential, and industrial uses) for ∼ 12 million people ^23^. In this study, water samples at different depths were collected from two sites (i.e., S4 and S7) at the SR from four seasons (i.e., April 2017, August 2017, November 2017, and January 2018) representing the vertical mixing (i.e., April and January) and thermal stratification seasons (i.e., August and November). Algae bloom and thermal stratification were occurring in the SR from mid-July 2017 to November 2017, which made a gradient of oxygen between the highly oxygenated surface water to hypoxia/ anoxia bottom water. However, the water columns were well-oxygenated in April 2017 and January 2018. Details on reservoir seasonal-vertical hydrodynamics of this sampling campaign were described elsewhere ^12^. Water samples were collected at the depth of 0.5 m, 10 m, 20 m, and 40 m from sites S7 and S4, and water samples at the depth of 53 m were only collected from site S4 **(Fig. S1**). Based on decreasing pattern of dissolved oxygen (DO) during stratified seasons, water columns were split into two zones: the shallow zone (0-10 m) with the average DOs higher than 3 mg/L, and the deep zone (20-53 m) with the average DOs lower than 3 mg/L. A total of 16 physico-chemical parameters and 22 micropollutant compounds were measured previously ^24^, from which the most frequently detected micropollutants (> 30%; 17 micropollutants) were included in further analysis (**Table S1**). Water samples (500 – 1,000 mL) were filtered through 3 μm filters and 0.22 μm filters to get size-fractionated sub-samples for DNA extraction. DNA retained onto 3 µm filters was treated as the PA fraction while DNA retrained on the 0.22 μm filters was treated as the FL fraction.

### 2.2 HT-qPCR and 16S rRNA gene amplicon sequencing

A total of 70 DNA samples from various depths (18 for April, August, and January and 16 for November) representing both PA and FL fractions were analyzed via HT-qPCR. HT-qPCR was performed using the Wafergen SmartChip Real-time PCR system with a total of 296 primer sets for 285 ARGs, 10 MGEs (eight transposase genes and two integrons), and 16S rRNA gene. HT-qPCR protocol was described in supplementary information S1. For the following analyses, otherwise mentioned, the abundance of ARGs or MGEs is always referred to as the normalized abundance (i.e., the ratio of ARG or MGE copy number to 16S rRNA gene copy number) ^25^. The ARGs with low normalized abundances (< 0.01% relative abundance of the total communities) were discarded. Also, three FL samples from January (at the depth of 20 m, 40 m, and 53 m from S4) and one FL sample from November (at the depth of 53 m from S4) were not included due to a low number of ARG occurrences (≤ 5; the mean number of ARGs per sample = 58.53). The absolute copy number of 16S rRNA genes was determined by using a LightCycler Roche 480 Real-time PCR system (Roche Inc., Basel, Switzerland). The detailed protocol was described in supplementary information S2.

The V4-V5 hypervariable regions of prokaryotic 16S rRNA genes were amplified using 515F (5’-GTG YCA GCM GCC GCG GTA-3’) and 907R (5’-CCG YCA ATT YMT TTR AGT TT-3’) and sequenced by using Illumina HiSeq 4000 sequencing platform. The raw reads generated were deposited in the NCBI short reads archive database under BioProject number PRJNA559031. Reads were clustered into operational taxonomic units (OTUs) with 97% sequence similarity. Taxonomic assignment was performed using the RDP classifier with SILVA database v132 ^14^. Rare taxa (relative abundance of less than 0.01% per sample and observed in less than 10 samples in PA and FL) fractions were excluded.

### 2.3 Null model analysis

Numerous studies applied the null model analysis to check whether the difference in microbial community composition and structure in habitat was associated with the variations of alpha-, gamma-diversity, or randomization ^26^. Here, a null model-based stochasticity ratio was used to explore the strength of stochastic processes in governing the assembly of PA and FL ARGs ^5^. This model firstly calculated the expected similarity community based on 1,000 randomizations of original ARG community data and the proportion of species occupancy was kept the same with the observed community. The relative importance of stochastic processes was then calculated by using the formula ^5^:

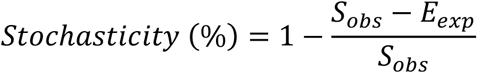

Where *S*_*obs*_ refers to the observed total similarity and *E*_*exp*_ refers to the mean of expected similarity from the null communities.

### 2.4 Particle-association niche (PAN) index

Previous studies indicated the particle-association niche (PAN) index can be used to characterize microbial lifestyle preferences (in PA and FL) ^16,19^. Here, we adopted it to define the lifestyle preference of ARGs. PAN index of each ARGs subtype was calculated using the below formula extracted from the R script provided by Salazar et al. (2015) ^16^:

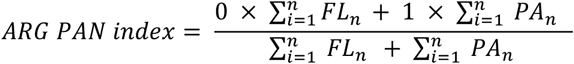

Where, FL refers to the abundance of ARG subtype in the FL fraction, PA refers to the abundance of ARG subtype in the PA fraction, and n refers to the number of samples. The ARG abundance in the FL fraction was given a weight of 0, while the ARG abundance in the PA fraction was given a weight of 1 creating a continuous index from 0 (strictly FL ARGs) to 1 (strictly PA ARGs). This index was then compared against a null distribution of the PAN index generated from 1,000 permutations ^16^. ARGs with significant indexes were defined as conservative ARGs (for PA or FL fractions), while those without non-significant indexes were defined as non-conservative ARGs (no lifestyle preferences). Besides ARGs, we employed this index for bacterial communities to explore the potential links between ARGs and bacteria with the same lifestyle. In addition, we calculated the PAN index for each ARG type (ARGs conferring resistance to the same antibiotics) and total ARGs.

### 2.5 Statistical analysis

To reveal the seasonal-vertical pattern of ARG communities in the SR, we performed multivariate analysis including permutational multivariate analysis of variance (PERMANOVA) and principal components analysis (PCA). Before performing PCA, the normalized abundance of ARGs was central log-ratio transformed ^27^. PERMANOVA was conducted with the ‘adonis’ function in the R package vegan ^28^. Correlation analysis was done by using Spearman correlation and the *P*-values were adjusted using the Benjamini-Hochberg method. The Significance test for two-group comparison was done by using the Wilcoxon test, while the significance test for multiple groups was done by using the Kruskal-Wallis test. Dunn test with Benjamini-Hochberg *P-value* correction was performed for pairwise comparison in multiple group comparisons.

The partial least squares path modeling (PLS-PM) ^29^ was used to explore the relationships between various factors and ARG abundances (in PA and FL fractions) or lifestyle transition (between PA and FL fractions as indicated by the PAN index). The hypothetical causation model was built by grouping variables based on their traits as follows: environmental variables (EVs), micropollutants (MPs), MGEs, bacterial biomass (16S rRNA gene abundance), and bacterial communities (BC). The initial model showed that temperature had distinct effects compared to other parameters in EVs. Therefore, it was separated as an independent variable. Moreover, to investigate the effects of the assembly of ARG communities on the lifestyle transition of ARGs, the stochasticity ratios between each pair of PA and FL ARG communities (ARGs stochasticity) were included in the PLS-PM model. In this model, BC, MGEs, and ARGs were expressed by using their PAN indexes. Only parameters with high correlations to its variable (loading ≥ 0.7) were included in the final PLS-PM model. The significance of the path coefficient and R^2^ in all PLS-PM was validated by using 1,000 bootstraps. In addition, the projection pursuit regression (PPR) based on MGEs was used to calculate the potential of HGT in PA and FL in the SR across seasons. The principle behind PPR is the projection of high-dimensional data into low-dimensional space to extract patterns that occurred within the data. This analysis was done with MATLAB 2013 ^30^.

Correlation-based networks were constructed to reveal the co-occurrence relationships among bacterial OTUs and ARGs in PA and FL fractions. Only ARGs and bacteria OTUs detected more than in 10 samples (∼30% of PA or FL samples) were included in the network analysis. The correlation cut-off was set to 0.6 with the adjusted *P*-value that was less than 0.01. The dissimilarity between PA and FL networks was calculated as proposed by Poisot et al. (2012).

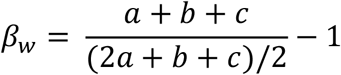

Where *β*_*w*_ is the dissimilarity between two networks, *a* is the number of shared edges, *b* is the number of unique edges for network one, and c is the number of unique edges to network two.

## Results

### 3.1 Seasonal-vertical variations of ARGs in PA and FL fractions in the SR

HT-qPCR was applied to check the abundance of various ARGs and MGEs in PA and FL fractions in the SR. The results indicated that a total of 133 ARG and 10 MGE subtypes were detected in PA and FL fractions. The mean normalized abundance of ARGs was significantly higher in the PA fraction (0.07 copies per 16S rRNA gene) than in the FL fraction (0.05) (Wilcoxon test, *P* < 0.05) (**Fig. 1A**). The mean absolute abundance of ARGs was 1.9 × 10^5^ copies/mL in the FL fraction and 5.9 × 10^5^ copies/mL in the PA fraction (Wilcoxon test, *P* < 0.1), respectively (**Fig. S2**). The normalized abundances of several ARG and MGE types, such as aminoglycoside, multidrug, chloramphenicol, sulfonamide resistance genes, and transposase genes, were significantly higher in the PA fraction than in the FL fraction (Wilcoxon test, *P* < 0.05) (**Fig. 1A**).

**Figure 1.**
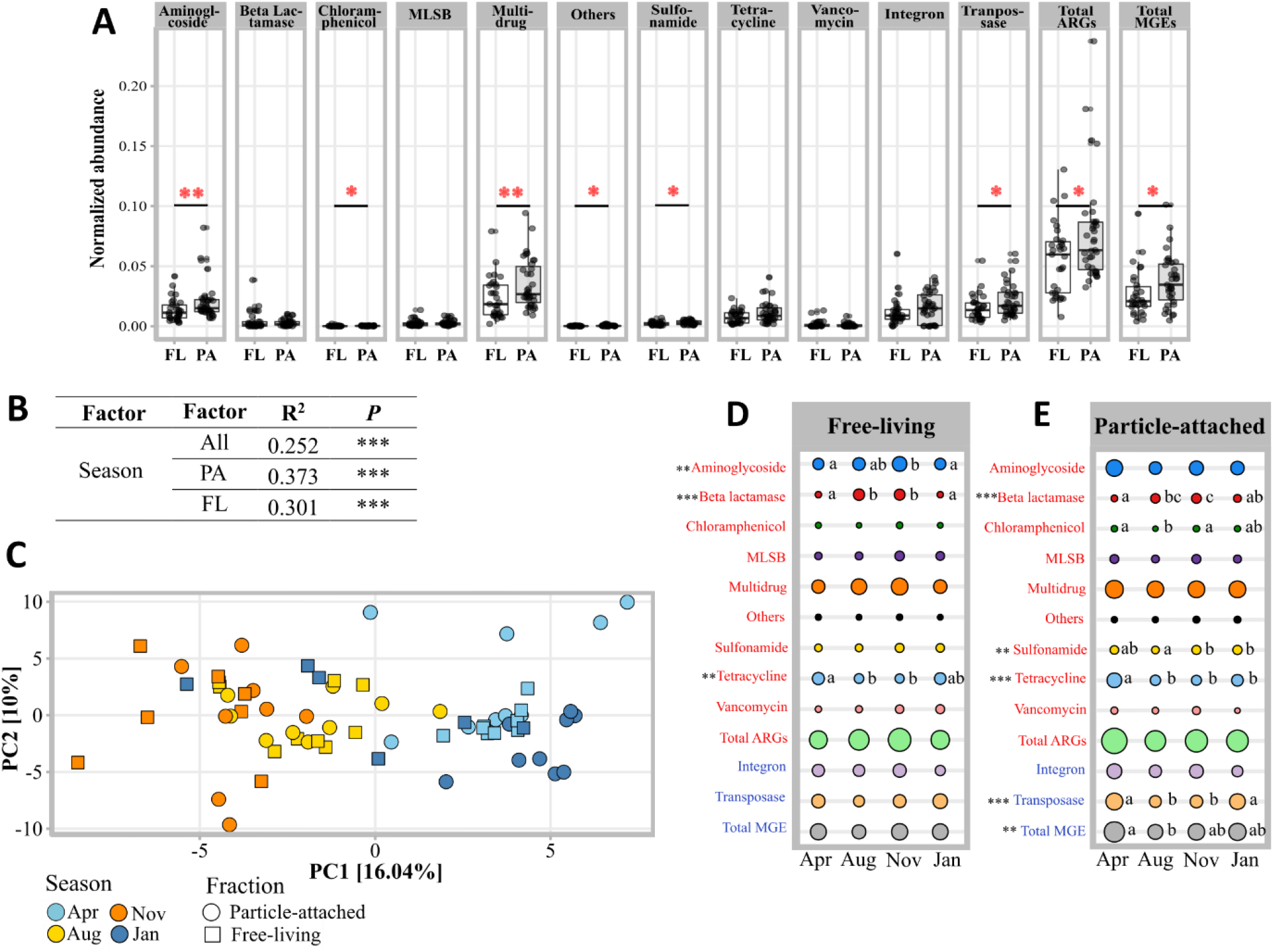
Pairwise comparison of the normalized abundance of ARG types, total ARGs, MGE types, and total MGEs between PA and FL lifestyles. Significant differences were tested by using the Wilcoxon test (* *P* < 0.05; ** *P* < 0.01; *** *P* < 0.001) (**A**), PERMANOVA analysis based on Bray-Curtis dissimilarity distance of ARGs with 9,999 permutations showing the variance of ARG communities in all (PA + FL), PA, and FL fractions explained by season (**B**). PCA plot showing the seasonal variation of the composition of ARG communities from PA and FL fractions in the SR (**C**). Seasonal differences of the normalized abundances of ARG types, total ARGs, MGEs (classified into integron and transposase), and total MGEs in PA (**D**) and FL fractions (**E**). Significant differences were tested by using the Kruskal test (*** *P* ≤ 0.001; ** *P* ≤ 0.01; * *P* ≤ 0.05). Seasons with different letters indicate a significant difference (Dunn test, *P* ≤ 0.05).

Seasonal and vertical variations of the PA and FL ARGs were also investigated. PERMANOVA results indicated that ARG communities in the SR were mainly subjected to seasonal variations (PA fraction: R^2^ = 0.373, *P* < 0.001; FL fraction: R^2^ = 0.301, *P* < 0.001) (**Fig. 1B**). Being consistent with PERMANOVA results, the PCA analysis revealed seasonal patterns for both PA and FL ARG communities (**Fig. 1C**). Interestingly, different ARG types in PA and FL fractions showed different seasonal patterns. Specifically, aminoglycoside resistance genes showed significant seasonal variations in FL fraction only (Kruskal-Wallis test, *P* < 0.01), while sulfonamide (*P* < 0.01) and chloramphenicol (*P* < 0.01) resistance genes showed significant seasonal differences in their normalized abundances in PA communities (**Fig. 1D & E**). Moreover, seasonal variations of transposase genes (*P* < 0.05) and the total MGEs (*P* < 0.01) were only occurring in PA communities.

The vertical stratification in SR water columns was pronounced during the stratified seasons (from August to November) as shown by the depletion of dissolved oxygen from surface to bottom water (**Fig. 2A)**. Being consistent with this pattern, PERMANOVA tests suggested that depth played a more important role in shaping the PA and FL ARG communities in stratified seasons (PA fraction: *P* < 0.05, FL fraction: *P* < 0.05 in August; FL fraction: *P* < 0.05 in November) compared to seasons with well-mixed water columns (PA and FL fractions: *P* > 0.05 in April and January) (**Fig. 2D**). Although ARG abundances in PA and FL fractions were comparable during the stratified period, the abundance of ARGs in both fractions was different between shallow and deep water (**Fig. 2B & Fig. S3**). For example, beta-lactamase resistance genes tended to be more abundant in shallow water in PA and FL fractions (Wilcoxon test, *P* < 0.05) (**Fig. 2C)**, while multidrug-, sulfonamide-, vancomycin-resistance genes and MGEs were more abundant in the deep water in PA and FL fractions (Wilcoxon test, *P* < 0.05). For seasons with well-mixed water columns (i.e., April and January), the vertical differences in the abundance of ARGs and MGEs were smaller.

**Figure 2.**
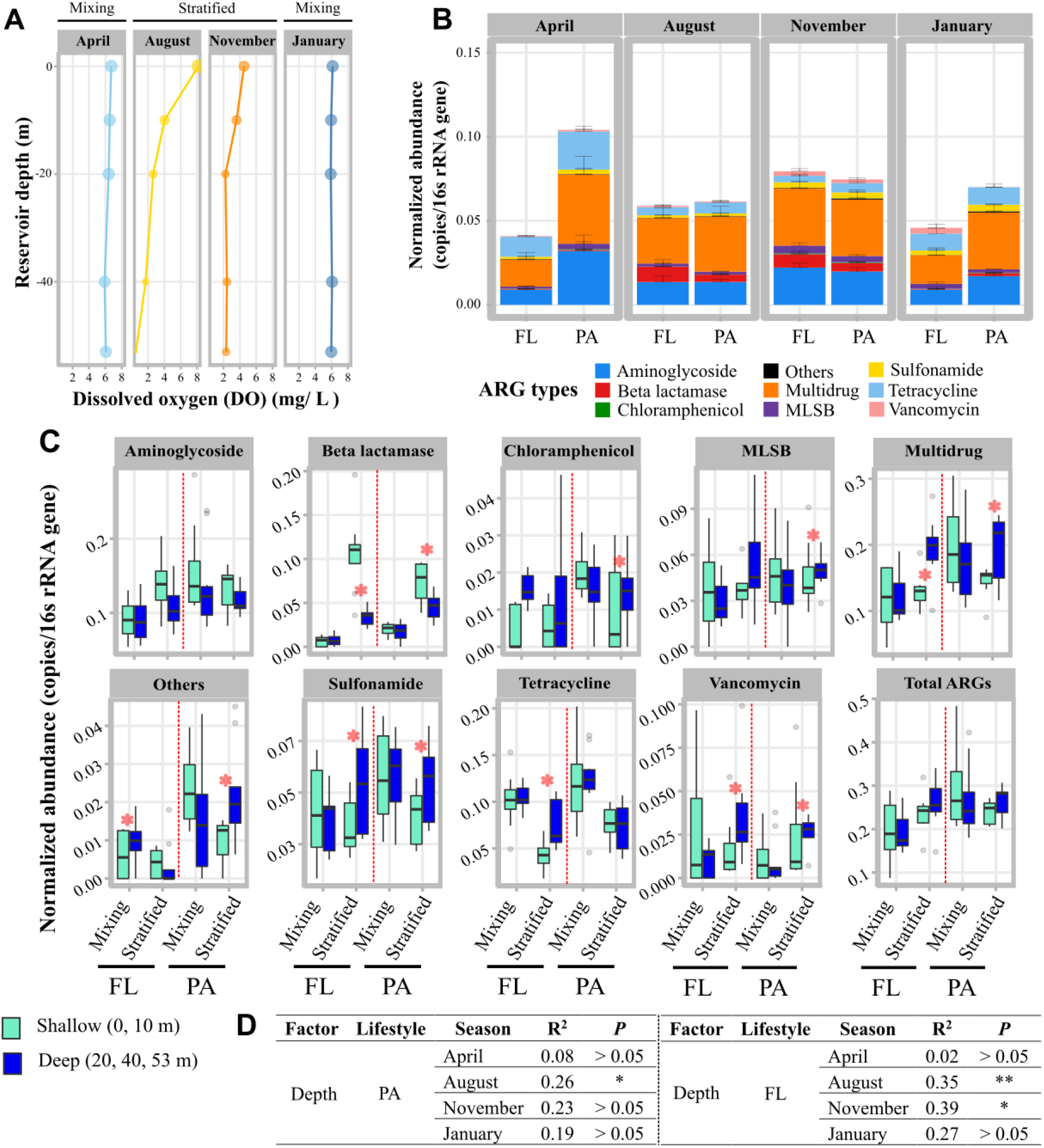
Vertical profile of dissolved oxygen (DO) in the water column of SR at four seasons (**A**), normalized abundances of ARG types including aminoglycosides, beta-lactamase, chloramphenicol, macrolide-lincosamide-streptogramin B (MLSB), multidrug, sulfonamide, tetracycline, vancomycin and others (i.e., other ARG types) at different seasons with error bars indicate standard errors of samples (**B**), and Pairwise comparisons between the normalized abundance of ARG types at the shallow and deep zones for PA and FL in seasons with well-mixed water columns (Mixing) and stratified seasons (Stratified) (Wilcoxon test, * *P* < 0.05) (**C**), PERMANOVA analysis based on Bray-Curtis dissimilarity matrix explaining the variances of ARG communities of PA and FL lifestyles explained by depth during stratified periods (i.e., August and November) and mixing periods (i.e., April and January) (* *P* < 0.05, ** *P* < 0.01) (**D**)

### 3.2 Transition of ARGs between PA and FL lifestyles among different hydrological seasons

The PAN index analysis indicated that there were 23 PA conservative ARGs, 16 FL conservative ARGs, and 94 ARGs with non-conservative lifestyles (no lifestyle preferences) (**Fig.3**). However, the lifestyles of these ARGs were not consistent among different seasons, suggesting these ARG subtypes did not strictly maintain one lifestyle across seasons in the SR. Nevertheless, there was a noticeable reduction of PA and FL conservative ARGs from seasons with well-mixed water columns (April: PA = 11, FL = 9; January: PA = 4, FL = 11) to stratified seasons (August: PA = 3, FL = 2; November: PA = 2, FL = 1) (**Fig. 3B, Table S2**). Furthermore, the total ARGs tended to be more abundant in the PA fraction (PAN index > 0.5) in the seasons with well-mixed water columns than in stratified seasons. Interestingly, the total ARGs became equally abundant in PA and FL fractions (PAN index ≈ 0.5) at stratified seasons (i.e., August and November) (*P* < 0.001). These results suggested a transition from PA lifestyles to FL lifestyles (**Fig. 3C, Fig. S3**).

**Figure 3.**
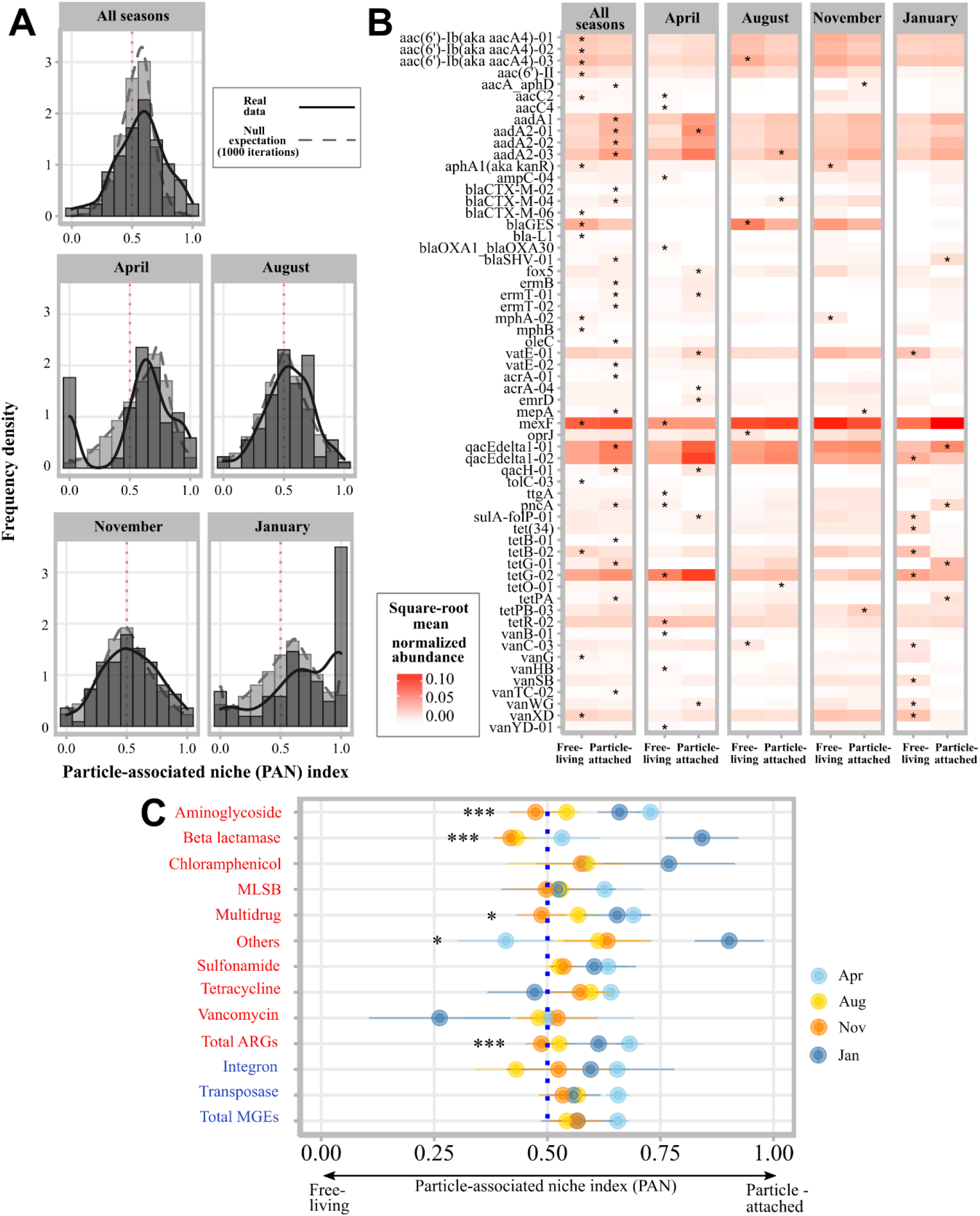
Histogram of the observed PAN index distribution for each ARG subtype compared to the expected PAN index from the null model (1,000 permutations) in the whole dataset and each season (**A**). Heatmap of ARG subtype with significant lifestyle (in PA or FL lifestyle) in all and individual season (**B**). The seasonal variations of the lifestyle shift (i.e., PAN index) of ARG types, total ARGs, MGE types, and total MGEs (Kruskal Wallis test: *** *P* ≤ 0.001; ** *P* ≤ 0.01; * *P* ≤ 0.05) (**C**).

### 3.3 Relative impacts of abiotic and biotic factors on ARGs in PA and FL fractions

PLS-PM was used to decipher the effects of abiotic and biotic factors regulating the abundances of ARGs in PA and FL fractions in the SR. The results indicated that MGEs were the main factor shaping the abundances of PA (total effect = 0.79) (**Fig. 4A & 4B**) and FL (total effect = 0.5) ARGs (**Fig. 4C & 4D**). Although the total effect of the bacterial communities on the PA ARG abundances was as high as MGEs (total effect = 0.74), the bacterial communities’ effects on the FL ARG abundances were half of the effects caused by MGEs (total effect = 0.26) (**Fig. 4B & 4D**). Environmental variables’ negative effects (total effect = -0.44) and MGEs’ positive effects were comparable in the FL fraction, while environmental variables had lower positive total effects than MGEs in the PA fraction (total effect = 0.38). It’s worth noting that micropollutants played a less important role in shaping the PA (total effect = 0.27) than FL (total effect = 0.37) ARG abundances. PLS-PM analysis also showed that MGEs were the main factor influencing the lifestyle shift of ARGs (total effect = 0.62), followed by the bacterial communities (total effect = 0.46) (**Fig 4E & 4F**). For abiotic factors, the environmental variables (e.g., DO, NO_2_-N, and DOP) had a similar impact on the lifestyle shift of ARGs as the bacterial communities (total effect = 0.41) (**Table S3**). Remarkably, the temperature had a negative total effect on the lifestyle shift of ARGs (total effect = -0.34).

**Figure 4.**
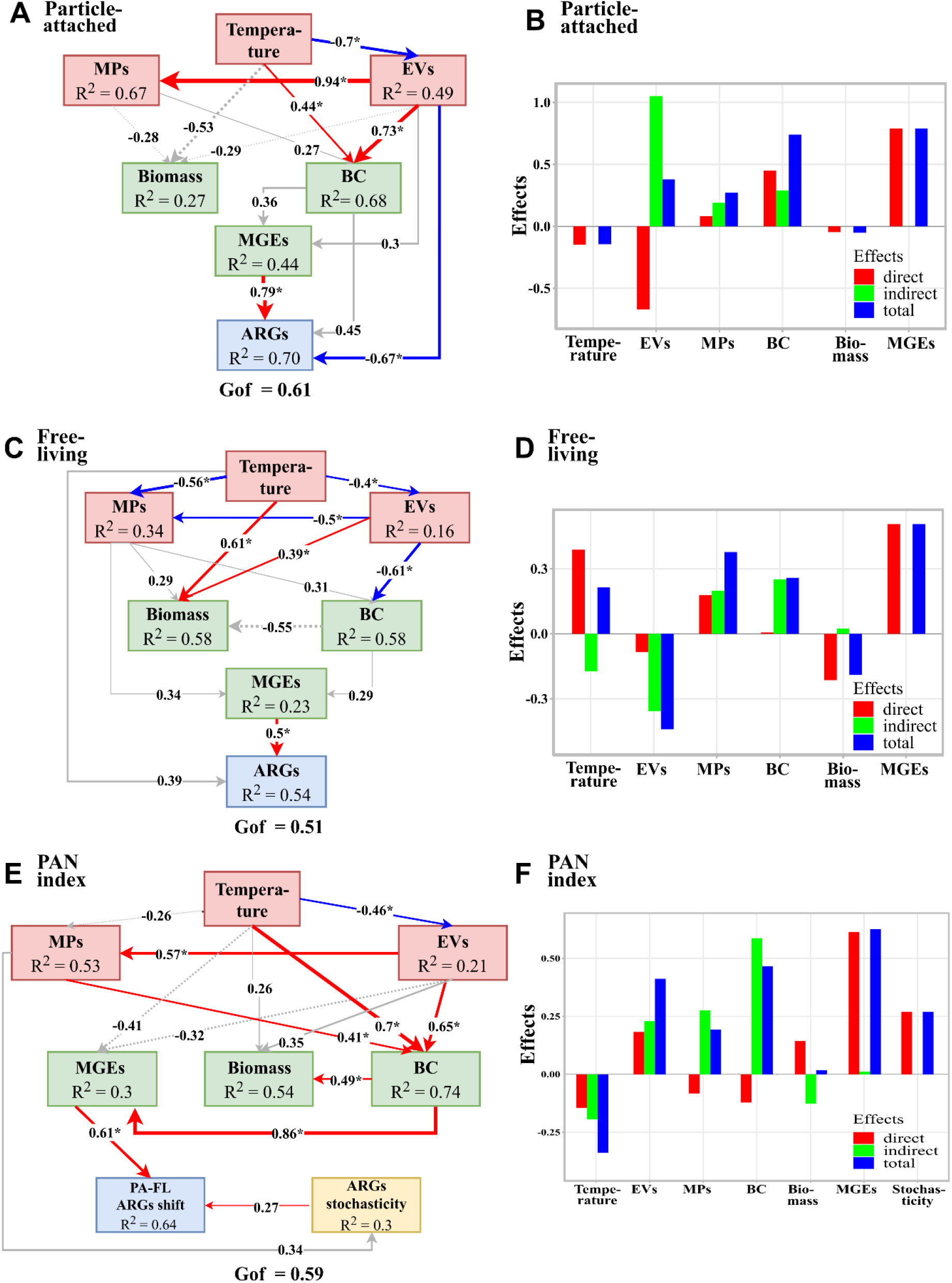
PLS-PM showing the effects of different factors on the total normalized abundance of ARGs in PA (**A**) and FL fractions (**C**), and lifestyle shift of ARGs (i.e., PAN index) (**E**). Variables included for the PA and FL models were bacterial communities (BC), micropollutants (MPs), environmental variables (EVs), MGEs, and temperature. For the PAN model, ARG community stochasticity was included, and the PAN index of bacterial communities (BC), FL bacterial abundance, and PAN index of MGEs were used. Dashed and solid lines represent the negative and positive paths. Significant paths are indicated by the asterisk (* *P* < 0.05) and colors (red: positive; blue: negative). The path coefficients (R^2^) and total standardized effects were calculated after 1,000 bootstraps. Models were assessed with the goodness of fit (GoF) values. Total standardized effects from each latent variable for PA ARGs (**B**), FL ARGs (**D**), and lifestyle shift of ARGs (**F**) were shown in the bar chart.

### 3.4 Co-occurrence associations between ARGs and bacterial taxa

A total of 409 and 402 ARG-bacterial OTU associations were found in PA and FL fractions, respectively. The high number of unique associations between ARGs and bacterial OTUs in each network (unique associations for PA = 336; unique associations for FL = 329) resulted in high dissimilarity (*β*_*w*_ = 0.82) between the two networks (**Fig. 5**). Interestingly, the shared ARG-bacterial OTU associations between PA and FL networks were mainly related to the non-conservative ARGs (≈ 66%) and FL conservative ARGs (≈ 34%), but not the PA conservative ARGs. Additionally, there was a high proportion of non-conservative ARGs in PA (24 of 37 ARGs and 300 of 409 associations) and FL (26 of 35 ARGs and 304 of 402 associations) networks. No significant differences were observed between the number of associations of these non-conservative ARGs and bacterial OTUs in the PA and FL networks (*P* > 0.05). However, some PA conservative ARGs had significantly higher associations with bacterial OTUs in the PA network compared to the ones in the FL network (*P* < 0.05). For example, one of the PA conservative ARGs, *ermT-02* (macrolide-lincosamide-streptogramin B (MLSB) resistance), had 16 associations in the PA network but 0 association in the FL network. The same pattern was found for the FL conservative ARGs in the FL network. For instance, aac(6’)-II (aminoglycoside) and *vanXD* (vancomycin), which belonged to FL conservative ARGs, had higher associations in the FL network (aac(6’)-II: 13; *vanXD*: 8) than those in the PA network (aac(6’)-II: 1; *vanXD*: 3). Furthermore, the number of associations between conservative ARGs and conservative bacteria was higher in the FL network (16 associations) than in the PA network (1 association).

**Fig. 5.**
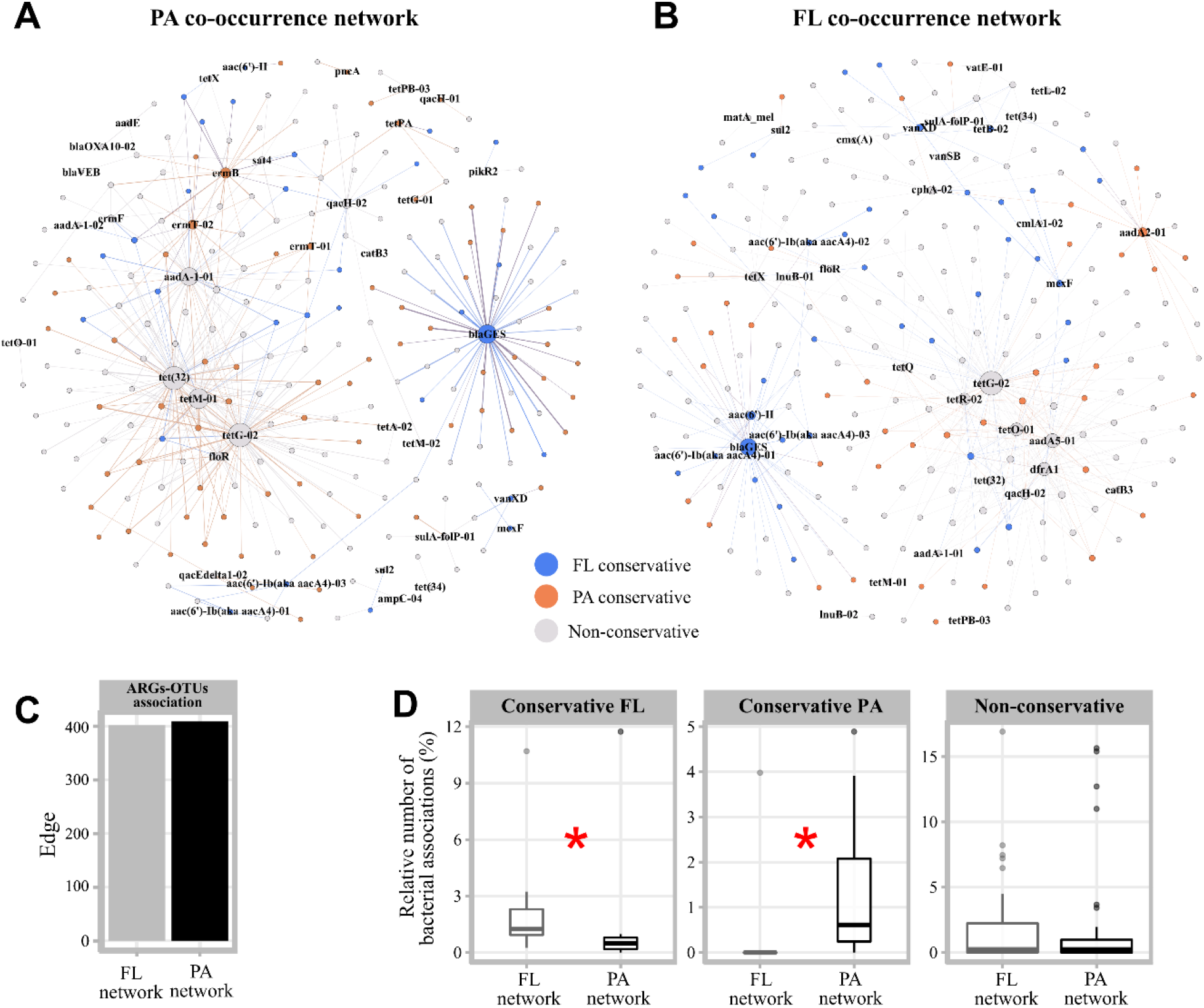
Associations between ARGs and bacterial OTUs in PA (**A**) and FL (**B**) networks, The number of ARGs-OTUs associations in the PA and FL network (**C**). A comparison of degree numbers between conservative and non-conservative ARGs in different lifestyles (**D**), Significant test was done using the Wilcoxon test (* *P* < 0.05).

## Discussion

Antibiotic resistance genes (ARGs) have been recognized as global emerging contaminants ^32^, and the pathways and mechanisms of ARG dissemination in deep freshwater reservoirs have not been investigated thoroughly, especially for PA and FL ARGs. Our results indicated that the seasonal change was the main driver of the ARG communities (in PA and FL) in the deep-water reservoir. Similar seasonal variations of ARGs were observed in different aquatic environments ^3,33,34^. However, the dynamics of PA and FL ARGs were controlled by different water physico-chemical parameters induced by the vertical mixing and thermal stratification in the SR. PLS-PM models indicated that the abundances of PA ARGs were mainly influenced by the nutrient loading such as NH_4_-N, DTP, and DOP, which were highest during April (the season with well-mixed water columns) (**Fig. S4**). Nutrients have been found to be correlated with the abundances of certain ARG types (e.g., tetracycline, MLSB, and sulfonamide) in a riverine ecosystem ^34^, which could be due to their function in enhancing cell metabolisms and activities ^35^. The particles containing ARGs in water columns may come from sediment resuspension during seasons with well-mixed water columns (i.e., January and April) ^12^ or upstream river deposits. Additionally, positive correlations between ARGs with different physico-chemical parameters, such as the relationship between sulfonamide resistance genes and NO_3_-N or soluble reactive phosphorus, the relationship between tetracycline resistance genes and DO, might be due to the seasonal variation of the microbial community carrying these ARGs ^36^.

In addition, the abundances of FL ARGs in hypoxia water columns during the stratified seasons were significantly higher than that of seasons with well-mixed water columns. We suggested that this phenomenon could be related to the seasonal dependent-lifestyle transition pattern of ARGs between the PA and FL lifestyles (see discussion below). Moreover, the positive correlations between certain FL ARGs (e.g., tetracycline and sulfonamide resistance genes) and nutrients indicated unique seasonal patterns which were similar to the ones for PA ARGs. This could be due to the detachment of ARG-carrying microbes from particles to live freely ^37^.

As a typical subtropical deep freshwater ecosystem, the SR experienced a seasonal vertical stratification water column starting from summer to fall ^12^. Similarly, we observed a vertical differentiation of ARGs profile from the surface water to bottom water (**Fig. 2C**). The increased abundance of multidrug-, sulfonamide-, and vancomycin-related ARGs towards hypoxic zones at the bottom of the SR might be due to the sinking and/or resuspension of ARGs containing particles from sediments ^38^. A previous study reported an increasing trend of tetracycline and integron abundance with depth in a subalpine lake during stratification periods ^38^. Suspended particles were known to be vectors for transporting surface microbiome to deep water in marine environments ^19^. Our results hereby suggested that ARGs might be transported with the same mechanism. Correspondingly, beta-lactamase resistance genes were higher in the shallow water of the SR compared to the bottom water (**Fig. 2C**). Its strong significant correlations with temperature and chlorophyll-a (**Fig. S5**) may indicate that the phytoplankton bloom might play an important role in beta-lactamase resistance gene dissemination ^39^. Overall, our results highlight that the reservoir undergoing seasonal vertical mixing and thermal stratification may regulate ARGs in PA and FL differently through the changes of abiotic (e.g., physico-chemical parameters) factors. Although we found a specific significant positive association between sulfonamide resistance genes and the concentration of sulfamethoxazole in PA and FL fractions (**Fig. S5**), the impact of micropollutants on the dissemination of whole ARGs may be insignificant due to their low concentrations in the SR ^14^.

Mounting evidence suggests that PA and FL fractions have distinct microbial communities in various aquatic environments ^16,19^. However, some recent evidence showed that PA and FL microbial communities were connected and could change lifestyles in response to environmental changes ^21,37,40^. Although a certain number of studies have explored PA and FL resistome in various environments ^10,41,42^, we first investigate the ARG dynamic of an aquatic system by taking into account the connectivity between PA and FL lifestyles. By using the PAN index as a proxy of lifestyle preference, we found that most ARGs had a non-conservative lifestyle in the SR. Despite that, several conservative ARG subtypes in both PA and FL fractions tended to have more associations with bacterial OTUs in their preferred lifestyle (**Fig. 5D)**. Additionally, we found associations between conservative ARGs and bacterial OTUs with the same lifestyle preference (one association in the PA network; 16 associations in the FL: network). This phenomenon may be due to a better adaptation of microbial communities carrying these ARGs in their preferred lifestyle than in the corresponding lifestyle ^16,43^. In this context, it also indicates that lifestyle could be an important factor determining the dissemination of specific ARGs.

Therefore, we incorporated the PAN index into PLS-PM to study the lifestyle transition between PA and FL ARGs. Our findings suggested that the temperature changes (total effects = -0.34) and DO (total effects = 0.4) might be critical factors indirectly driving the transition from PA to FL lifestyle by influencing the lifestyle transition of MGEs and bacteria (**Fig. 4E & 4F**). During stratified seasons, the SR experienced phytoplankton blooms (chlorophyll-a > 10 µg/L) and vertical stratification leading to hypoxia conditions in the deep layer of water columns ^12^. The micro-patches with low oxygen levels in particles have been found to be the critical factor regulating the PA microbial communities ^37,44^. Moreover, hypoxia conditions reduced the oxygen diffusion within the particles more than under the ambient oxygen conditions ^45^. Therefore, we argue that microbial taxa carrying ARGs and/or MGEs may shift the lifestyle (i.e., from PA to FL lifestyle) timely to maintain their metabolisms ^46^. Alternatively, the transition of ARGs between the PA and FL lifestyles could be related to microbial responses to the increasing dissolved organic carbon produced during phytoplankton blooms ^47^. A previous study found that the extensive bacteria exchange between attached and free-living communities during phytoplankton bloom could lead to an increase in microbial community similarity between PA and FL fractions ^21^. However, we are also aware that this phenomenon might be due to more complex abiotic and biotic interactions. Thereby, a controlled experiment is needed to further evaluate the factors contributing to PA and FL ARGs community transition. In our study, some evidence indicated that the PA rather than FL fractions may be more suitable micro-environments for ARG dissemination. Firstly, PLS-PM suggested that MGEs had a higher total effect on the abundance of ARGs in the PA fraction than that in the FL fraction (**Fig. 4**). In addition, the Spearman correlation analysis revealed a higher number of significant positive correlations between ARG and MGE types in the PA fraction (38) than in the FL fraction (28) (**Fig. S6**). Secondly, PPR analysis showed a higher potential of HGT rate in the PA fraction than the FL fraction across seasons (**Fig. S7**). Thirdly, the null model analysis indicated that the assembly of PA ARG communities had significantly higher stochasticity than that of FL ARG communities (**Fig. S8)**. These findings are consistent with the report of Yu et al. (2021) demonstrating that the PA ARG communities exhibit higher stochasticity than the FL ARG communities in the surface waters in the Yellow River ^42^. This phenomenon suggests that a higher probability of HGT occurred in the PA than in the FL fraction ^5^. Furthermore, the higher abundances of ARGs and MGEs in the PA than in the FL fraction (**Fig. 1A**) may as well be more favorable to promote ARG exchange in the PA fraction ^48,49^ due to the limited space and close contact between cells on particle ^48^. A shotgun metagenomics-based study on the marine environment reported that the higher abundances of ARGs and MGEs were responsible for social interactions and cell-to-cell transfer in the PA fraction rather than the FL fraction ^50^. This indicated that HGT may be a more common life strategy of microorganisms living on suspended particles. Intriguingly, the transition of ARGs from PA to FL fraction during stratified seasons might not only lead to an increase in the ARGs abundances (**Fig. S3**) but also enhance the HGT probability in FL fraction (**Fig. S9**).

In this study, we demonstrated that the size-fractionation of ARGs into PA and FL ARGs could be useful to model the lifestyle transition pattern of ARGs in a freshwater reservoir (**Fig. 6**). Such knowledge would be valuable in improving ARG pollution management in the environment. For example, the FL ARGs were more difficult to settle down during wastewater treatment plant (WWTP) processes ^41^ due to the lighter masses of their hosts compared to PA ARGs. Thus, controlling the factors leading to ARG desorption from the PA and FL fraction might minimize the ARG spread, especially in drinking water treatment plants and WWTPs ^41,51^.

**Figure 6.**
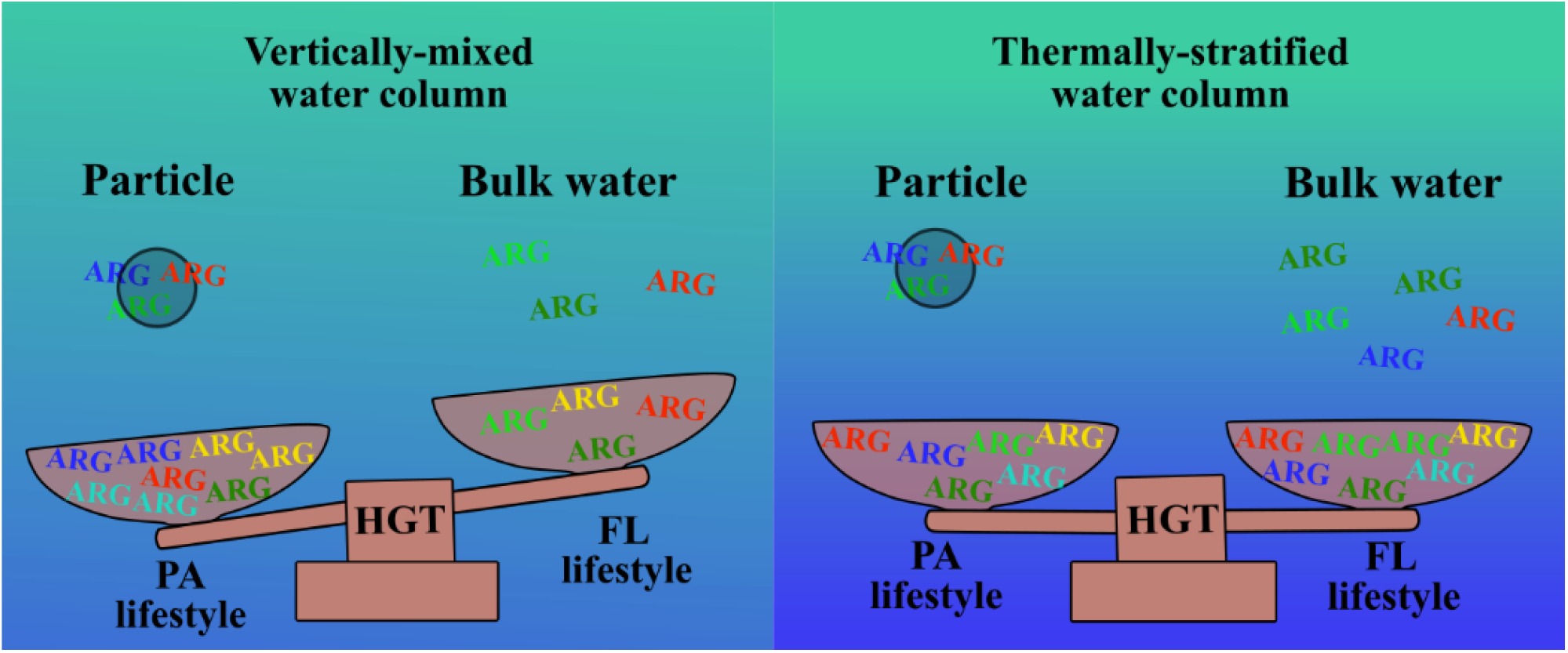
Conceptual diagram of inter-lifestyle ARGs transition (between PA and FL) under the influence of vertically-mixed and thermally-stratified water columns in the deep freshwater reservoir. The changes in physico-chemical parameters of the water column indirectly modulate ARGs transition through MGEs (HGT).

## Conclusion

We studied the influences of the seasonal hydrological dynamics (i.e., vertical mixing and thermal stratification) on the ARG profiles not only in both PA and FL fractions but also in their transition between lifestyles (e.g., from PA to FL lifestyle) in a deep freshwater reservoir. The main conclusions are as follows:

1. PA and FL ARGs responded differently to the changes in physico-chemical parameters induced by the seasonal-hydrological pattern (i.e., vertical mixing vs. thermal stratification). The abundance of PA ARGs increased in the mixing period characterized by high nutrient loads, while the abundance of FL ARGs increased in the stratified period characterized by severe DO depletion in water columns.
2. The seasonal-hydrological pattern shaped the transition of ARGs between the PA and FL lifestyles indirectly through the changes in water temperature and DO.
3. Despite a larger proportion of non-conservative ARGs, a fraction of conservative ARGs were found to be more dominant within their lifestyle network. Additionally, there were more ARG-bacteria associations with the same lifestyle preference in the FL than the PA network.
4. Although the HGT was the main mechanism driving the dissemination of ARGs in PA and FL fractions as well as between the PA and FL fractions, a higher potential for HGT was found in the PA than FL fractions.

## CRediT author statement

**Bob Adyari**: Conceptualization, Investigation, Methodology, Formal analysis, Software, Writing - original draft, Visualization. **Liyuan Hou**: Formal analysis, Validation, Writing - review & editing. **Lanping Zhang**: Investigation, Methodology, Data curation, Software. **Nengwang Chen**: Investigation, Methodology, Resources, Funding acquisition. **Feng Ju:** Methodology, Writing - review & editing. **Longji Zhu:** Methodology, Resources. **Chang-Ping Yu**: Resources, Supervision, Funding acquisition. **Anyi Hu**: Conceptualization, Data curation, Software, Validation, Writing - review & editing, Supervision, Project administration, Funding acquisition.

## Acknowledgement

This work was supported by the National Natural Science Foundation of China (U1805244 and 31870475), the CAS Youth Interdisciplinary Team (JCTD-2021-13), and the Youth Innovation Project of Xiamen (3502Z20206093). BA was supported by the CAS-TWAS president PhD fellowship programme.

## Notes

### Competing Interest Statement

The authors have declared no competing interest.

