## Supplemental Figures for "Hydrological dynamics drive the transition of antibiotic resistance genes between particle-attached and free-living lifestyles in a deep freshwater reservoir"

***Running title:*** *Transition of ARGs between PA and FL*

Bob Adyari^1,2,3,4^, Liyuan Hou^5^, Lanping Zhang^1,2,3^, Nengwang Chen^6^,

Feng Ju^7^, Longji Zhu^1^, Chang-Ping Yu^1,8^, Anyi Hu^1,2,3*^

1. CAS Key Laboratory of Urban pollutant Conversion, Institute of Urban Environment, Chinese Academy of Sciences, Xiamen 361021, China
2. University of Chinese Academy of Sciences, Beijing 100049, China
3. Fujian Key Laboratory of Watershed Ecology, Institute of Urban Environment, Chinese Academy of Sciences, Xiamen 361021, China
4. Department of Environmental Engineering, Universitas Pertamina, Jakarta 12220, Indonesia
5. Department of [Civil and Environmental Engineering](https://cee.usu.edu/), Utah state university, Utah, UT 84322, USA
6. Fujian Provincial Key Laboratory for Coastal Ecology and Environmental Studies, College of the Environment and Ecology, Xiamen University, Xiamen 361005, China
7. Key Laboratory of Coastal Environment and Resources of Zhejiang Province, School of Engineering, Westlake University, Hangzhou 310024, China.
8. Graduate Institute of Environmental Engineering, National Taiwan University, Taipei 106, Taiwan

***Correspondence to:**

**Supplementary information**

**Text S1: HT-qPCR protocol for 296 primer sets**

The total volume of the PCR reaction was 100 nL and the PCR reaction cycles were as follows: initial denaturation at 95°C for 10 mins, followed by 40 cycles of denaturation at 95°C for 30 s and annealing at 60°C for 30 s ^1,2^. All qPCR reactions were performed in triplicates with negative controls. Data were analyzed with SmartChip qPCR software (V2.7.0.1) and reactions with multiple peaks and amplification efficiency beyond 90-110% were discarded for each primer set ^1^. Cycle threshold (Ct) was set to 31 as the standard detection limit for qPCR and only samples with two qualified replicates were processed further. Copy numbers of ARGs were calculated by the following formula: gene copy number = 10^((31-Ct)/(10/3))^.

**Test S2: Real-time qPCR protocol for calculating absolute copy number of 16S rRNA**

A 20 µL of PCR reactions, containing 10 µL LightCycler 480 SYBR Green I Master Mix (2×) (Applied Bio-systems, CA, USA), 0.8 µL of each primer (0.4 µM), 0.5 mg/mL bovine serum albumin (Sigma, Steinheim, Germany), 2 μL of DNA template and 5.9 µL nuclease-free PCR grade water. The amplification routines were: 95°C for 10 mins, followed by 40 cycles of denaturation at 95°C for 30 s and annealing at 60°C for another 30 s, and extension at 72°C for 15 s. The standard curve was generated by using a tenfold serial dilution of 16S rRNA gene inserted plasmid ^2^

**Table S1.** The list of sixteen physico-chemicals and seventeen micropollutants measured in SR

(In excel file)

**Table S2.** Degree numbers of ARGs and bacterial OTUs of PA and FL co-occurrence networks.

(In excel file)

**Table S3.** PLS-PM models for PA, FL and particle-association niche (PAN) index.

(In excel file)

**Table S4.** Edges between conservative OTUs or ARGs (significant PAN index) in PA and FL co-occurrence networks.

(In excel file)

**Table S5**. Edges of PA and FL co-occurrence networks.

(In excel file)

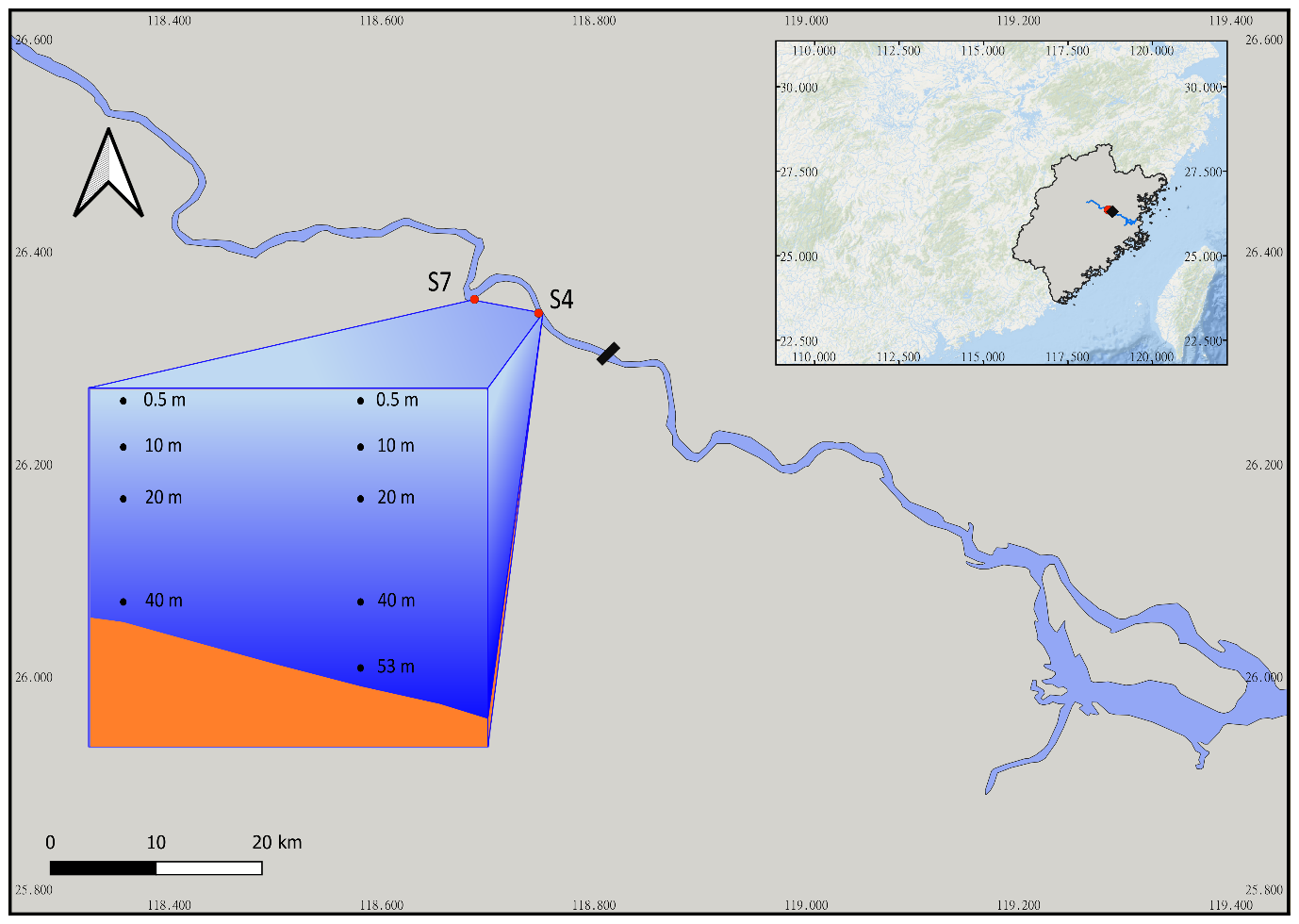

**Fig. S1.** The map of Shuikou Reservoir (SR) in Minjiang River revealing the location of sampling sites and the water column at S4 (0.5 m, 10 m, 20 m, 40 m, and 53 m below the water surface) and S7 (0.5 m, 10 m, 20 m, and 40 m below the water surface).

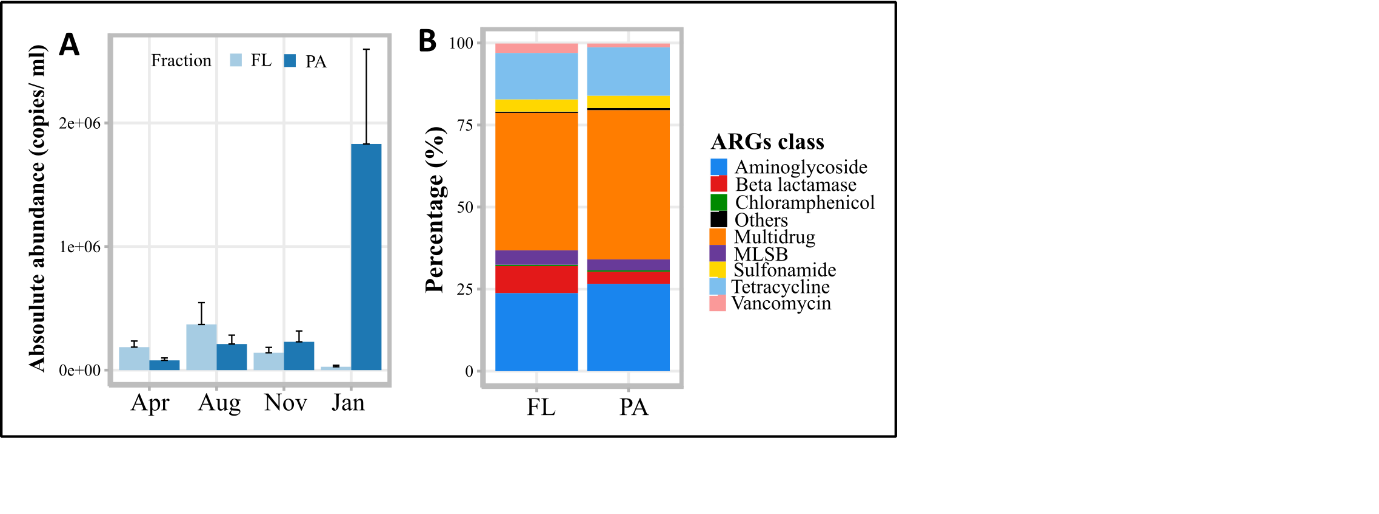

**Fig. S2.** The total absolute abundance of ARGs in PA and FL lifestyles in the SR at different seasons (**A**), and the composition of PA and FL ARG types (**B**).

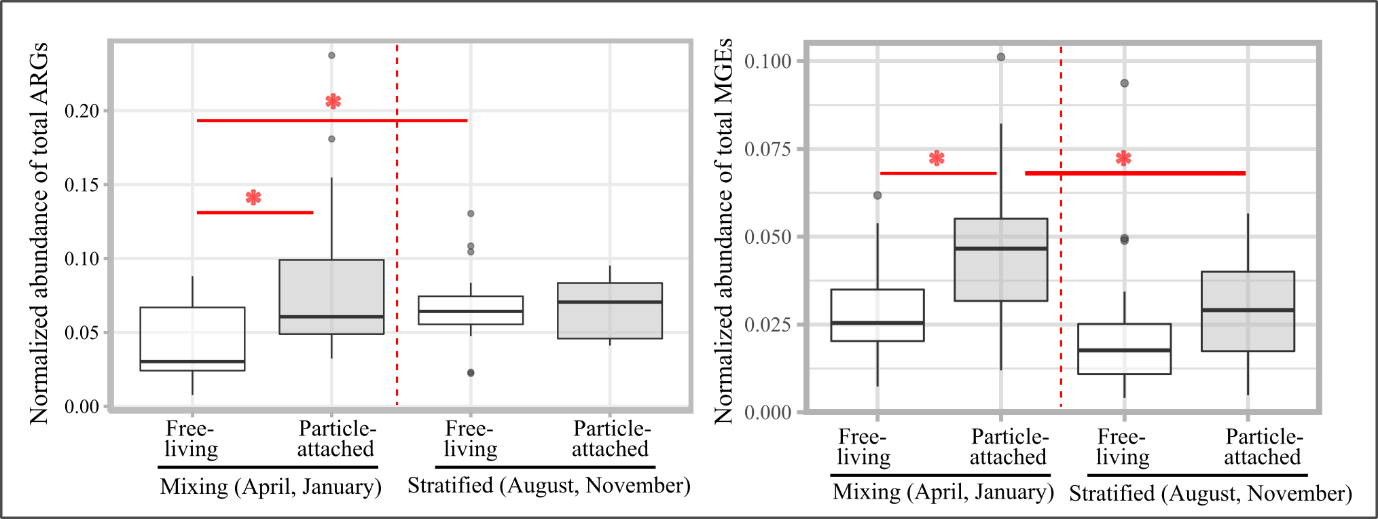

**Figure S3.** Comparison of normalized abundance of total ARGs and MGEs between seasons with well-mixed water columns (i.e, April and January) and stratified seasons (i.e, August and November) in PA and FL fractions. Significance differences were tested by using Wilcoxon test (* *P* < 0.05; ** *P* < 0.01; *** *P* < 0.001).

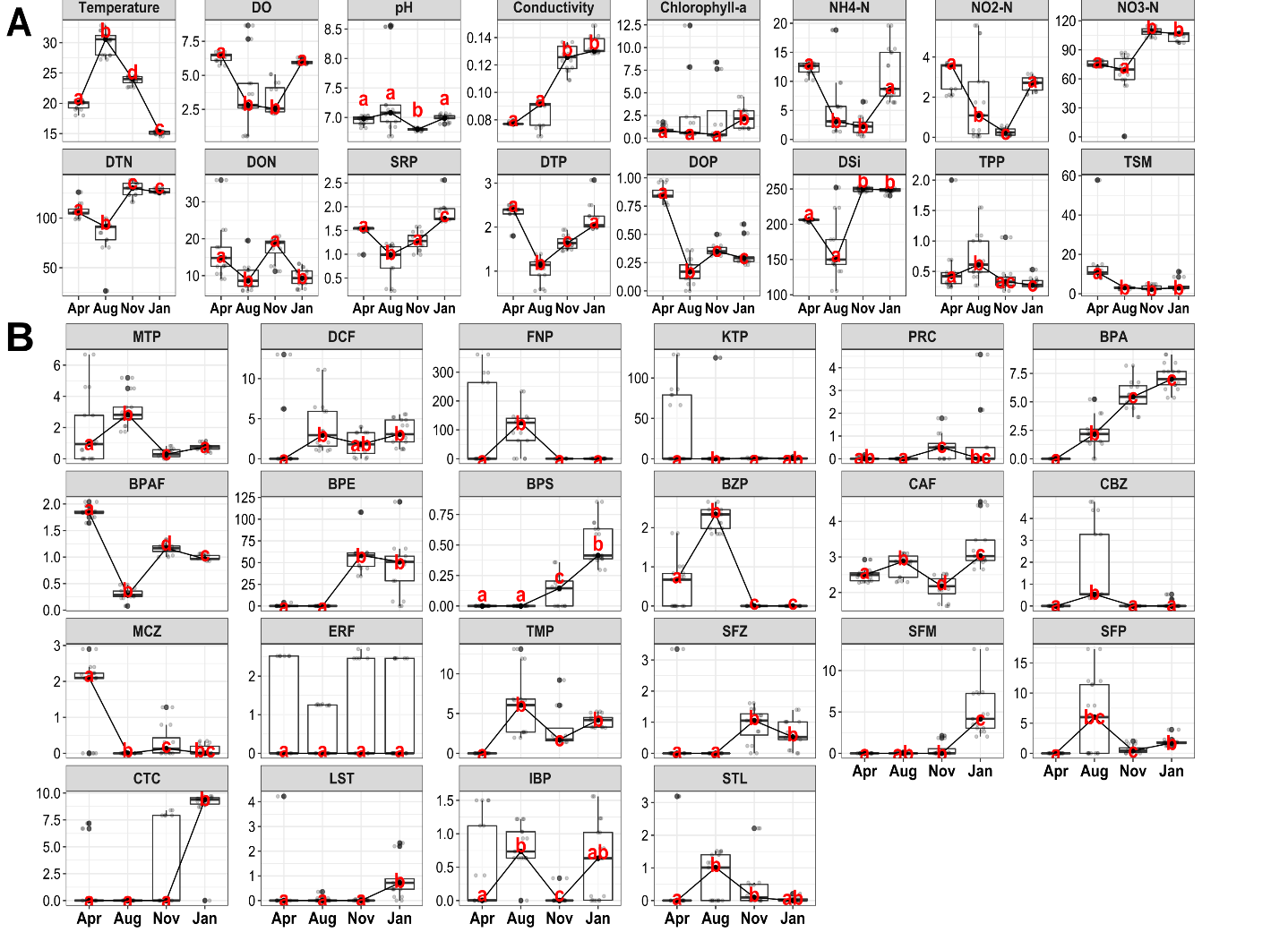

**Fig. S4.** Seasonal variation of physico-chemical (A) and micropollutants (B) in the SR. Significant differences were tested by using the Kruskal-Wallis test, followed by the pairwise Dunn test with Benjamini-Hochberg correction. Seasons without shared letters indicated significant differences (Dunn test, *P* < 0.05)

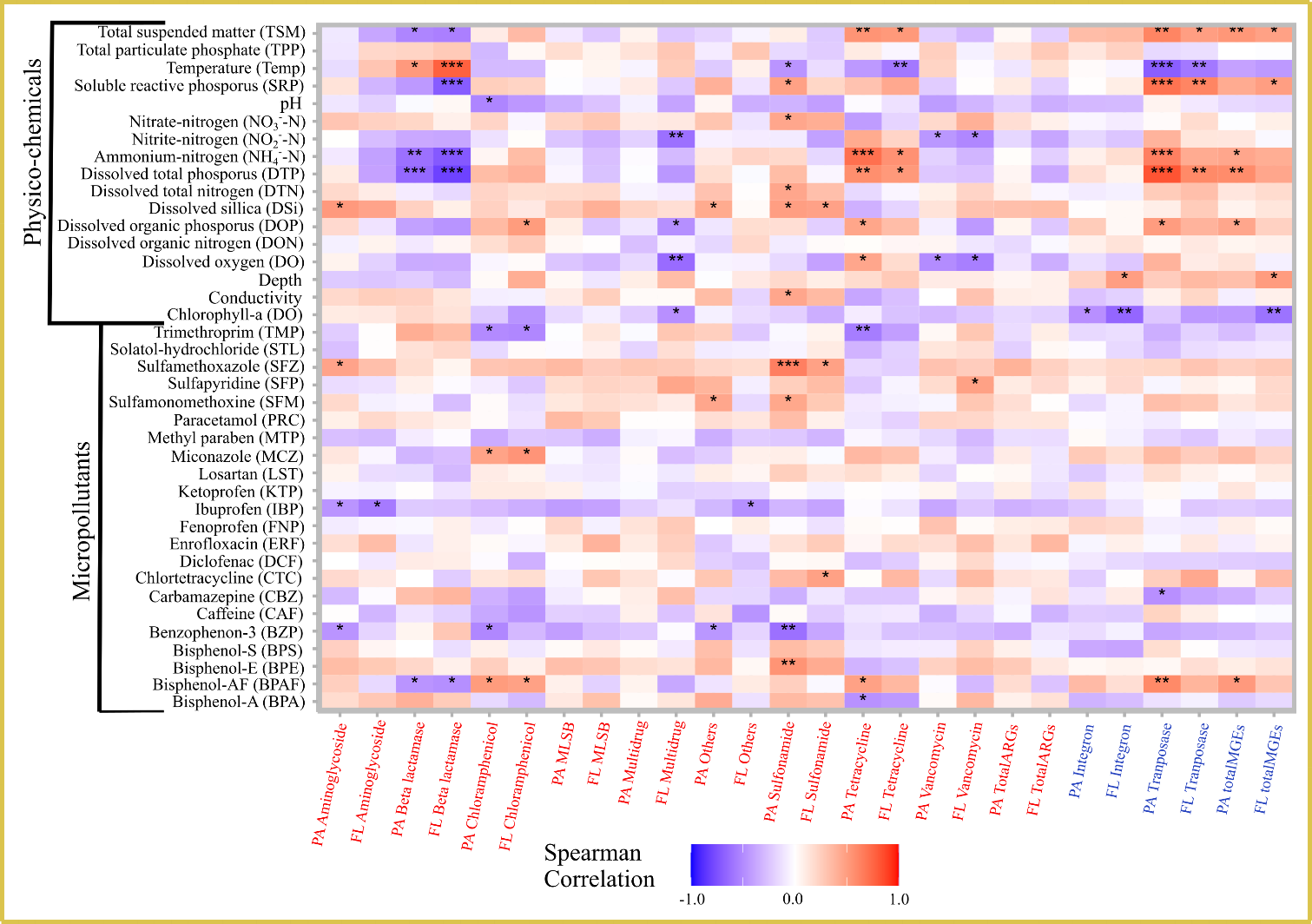

**Fig. S5.** A heatmap of the Spearman correlation among environmental variables (physico-chemicals and micropollutants), ARG, and MGE types in PA and FL lifestyles. Statistic significances were corrected further by using the Benjamini-Hochberg method (* *P* < 0.05; ** *P* < 0.01; *** *P* < 0.001).

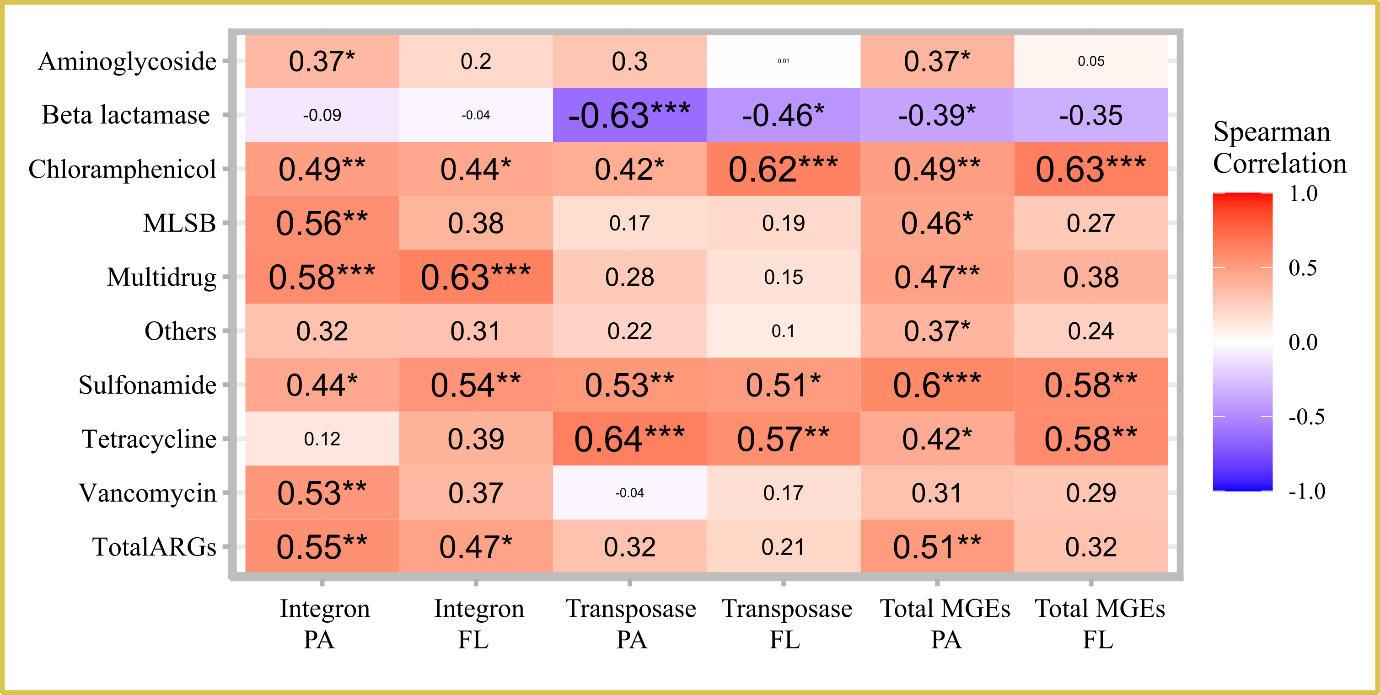

**Fig. S6.** A heatmap of Spearman correlations between ARG types in PA and FL lifestyles. The sizes of the values are proportional to the strength of the correlation. Statistic significances were further corrected by using the Benjamini-Hochberg method (* *P* < 0.05; ** *P* < 0.01; *** *P* < 0.001).

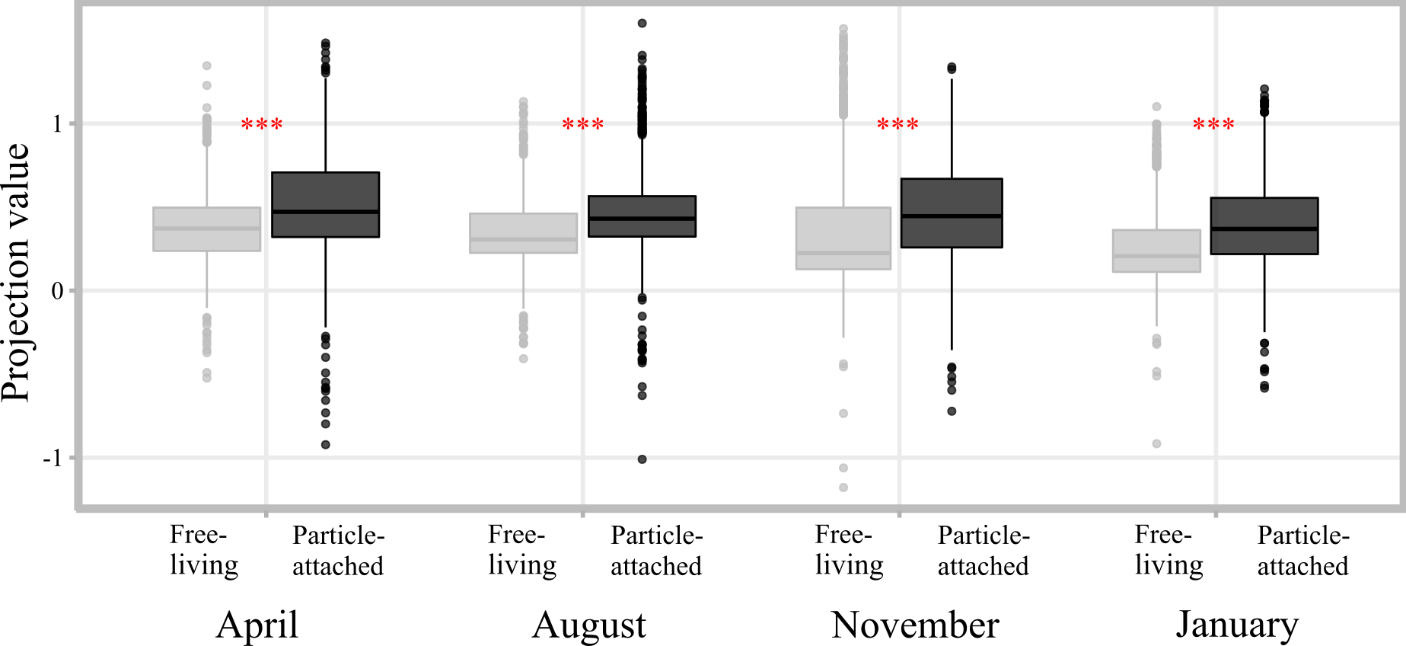

**Fig. S7.** Projection values of the HGT in different lifestyles and seasons in the SR based on all detected MGEs. The projection values are based on 100 times randomized trials. Significance differences were tested by using the Wilcoxon test (*** *P* < 0.001).

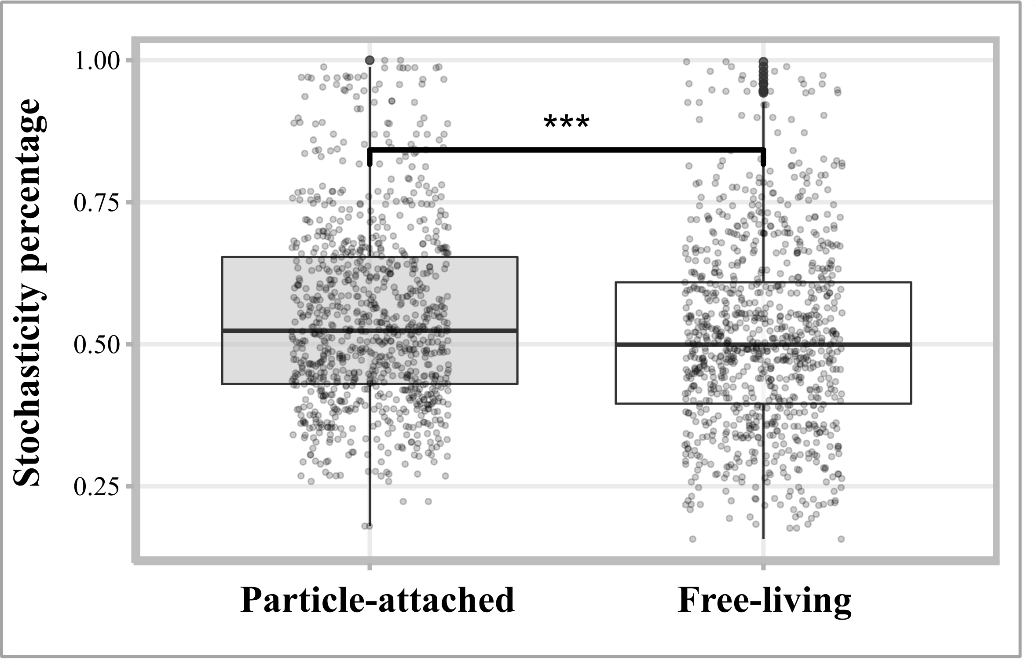

**Fig S8.** Stochasticity percentage representing the degree of stochasticity governing in PA and FL ARG communities. The significance test was conducted using the Wilcoxon test (*** *P* < 0.001).

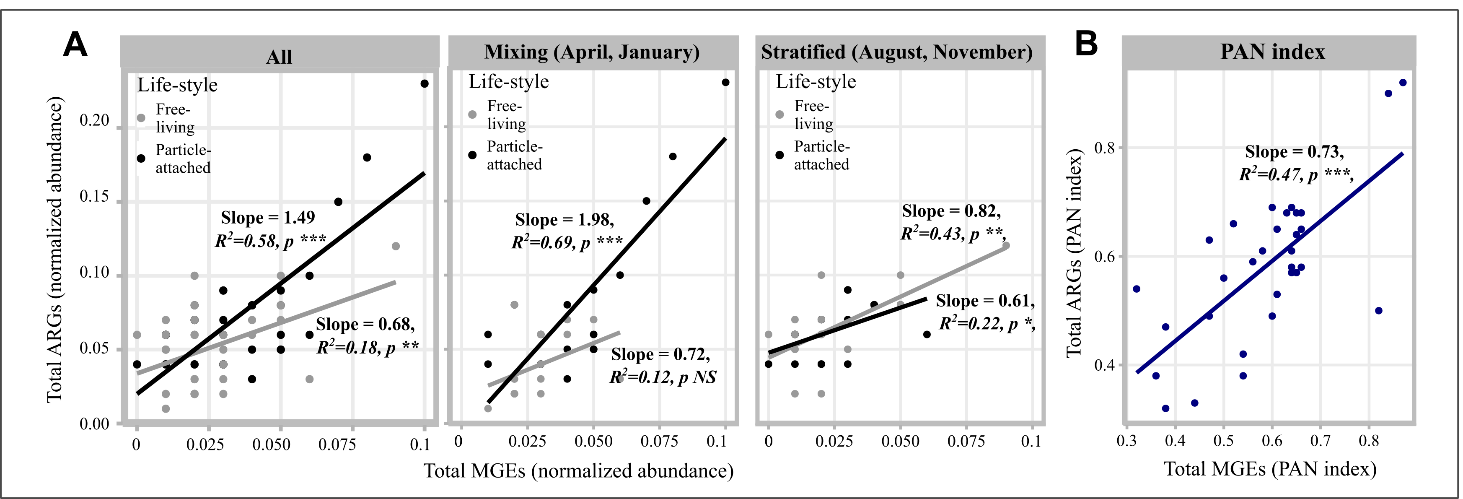

**Figure S9.** Relationship between normalized abundance of total MGEs and total ARGs in PA and FL (**A**) fractions and between PAN indexes of total MGEs and total ARGs (**B**) (*** *P* ≤ 0.001; ** *P* ≤ 0.01; * *P* ≤ 0.05, NS non-significant) (**C**).
